# Psilocybin induces rapid and persistent growth of dendritic spines in frontal cortex *in vivo*

**DOI:** 10.1101/2021.02.17.431629

**Authors:** Ling-Xiao Shao, Clara Liao, Ian Gregg, Pasha A. Davoudian, Neil K. Savalia, Kristina Delagarza, Alex C. Kwan

## Abstract

Psilocybin is a serotonergic psychedelic with untapped therapeutic potential. There are hints that the use of psychedelics can produce neural adaptations, although the extent and time scale of the impact in a mammalian brain are unknown. In this study, we used chronic two-photon microscopy to image longitudinally the apical dendritic spines of layer 5 pyramidal neurons in the mouse medial frontal cortex. We found that a single dose of psilocybin led to ∼10% increases in spine size and density, driven by an elevated spine formation rate. The structural remodeling occurred quickly within 24 hours and was persistent 1 month later. Psilocybin also ameliorated stress-related behavioral deficit and elevated excitatory neurotransmission. Overall, the results demonstrate that psilocybin-evoked synaptic rewiring in the cortex is fast and enduring, potentially providing a structural trace for long-term integration of experiences and lasting beneficial actions.

## Introduction

Serotonergic psychedelics are compounds that produce an atypical state of consciousness characterized by altered perception, cognition, and mood. It has long been recognized that these compounds may have therapeutic potential for neuropsychiatric disorders including depression, obsessive-compulsive disorder, and addiction (Nichols, 2016; Vollenweider and Preller, 2020). Among serotonergic psychedelics, psilocybin is recently shown to relieve depression symptoms rapidly and with sustained benefits for several months (Carhart-Harris et al., 2016; Davis et al., 2020; Griffiths et al., 2016; Ross et al., 2016). This progress led to a ‘Breakthrough Therapy’ designation by the FDA in 2019 and the initiation of multi-site clinical trials to test psilocybin as treatment for major depressive disorder.

It is well established that structural neuroplasticity in the frontal cortex is key to the action of antidepressants. Synaptic atrophy is found in the prefrontal cortex of patients with depression (Drevets et al., 1997; Holmes et al., 2019). Likewise, synaptic deficiencies including loss of dendritic arborization, reduced spine density, and damped neurotransmission are present in the frontal cortex of rodent chronic stress models (Cook and Wellman, 2004; Liston et al., 2006; Radley et al., 2004; Yuen et al., 2012). By contrast, compounds with fast-acting antidepressant effects promote structural plasticity to reverse the synaptic deficits caused by chronic stress (Duman and Aghajanian, 2012). For instance, a single dose of ketamine leads to higher spine density in the medial frontal cortex of rodents (Li et al., 2010), which is due to an increase in spine formation rate (Moda-Sava et al., 2019; Phoumthipphavong et al., 2016), likely involving elevated calcium signaling in the dendritic compartment (Ali et al., 2020a).

What is the current evidence that serotonergic psychedelics such as psilocybin can alter synaptic architecture? A few studies have shown that the expressions of genes involved in synaptic plasticity are elevated after administration of serotonergic psychedelics in rats (Nichols and Sanders-Bush, 2002; Vaidya et al., 1997). In neuronal cultures, bath application of serotonergic psychedelics induces transient increases in spine size (Jones et al., 2009) and proliferation of dendritic branches (Ly et al., 2018; Ly et al., 2020). A recent study showed that an analogue of ibogaine, a psychedelic with differing molecular targets from psilocybin, increases spine formation rate in mice (Cameron et al., 2020). Finally, in the pig, psilocybin administration was associated with higher binding of a presynaptic protein tracer in positron emission tomography (Raval et al., 2021). Although these studies provided clues linking serotonergic psychedelics to structural and functional neuroplasticity, significant gaps remain. In particular, there has been no direct demonstration of psilocybin-induced structural plasticity at cellular resolution in a mammalian brain. Importantly, the time scale in which such synaptic rewiring may occur *in vivo* is unknown.

## Results

### A single dose of psilocybin leads to long-lasting increases in spine density and spine head width in the mouse medial frontal cortex

To test the potency and dose dependence of psilocybin in mice, we measured the head-twitch response, a classic assay for characterizing psychedelic compounds in rodents. We observed that mice would exhibit high-frequency headshakes intermittently after administration of psilocybin (**Supplementary Video 1**). We characterized 82 C57BL/6J mice including 41 males and 41 females with 5 doses of psilocybin (0, 0.25, 0.5, 1, 2 mg/kg, i.p.; range = 7-10 per sex per dose). A sharp rise of elicited head-twitch responses occurred at 1 mg/kg (**Figure 1A**), consistent with prior reports (Halberstadt et al., 2011; Sherwood et al., 2020). Thus, we chose to use 1 mg/kg – the inflection point of the dose-dependence curve – to assess psilocybin’s effect on structural plasticity. At this dose, the rate of head-twitch responses peaked at 6–8 minutes after administration and then gradually declined until they ceased at about 2 hr (**Figure 1B**).

**Figure 1.**
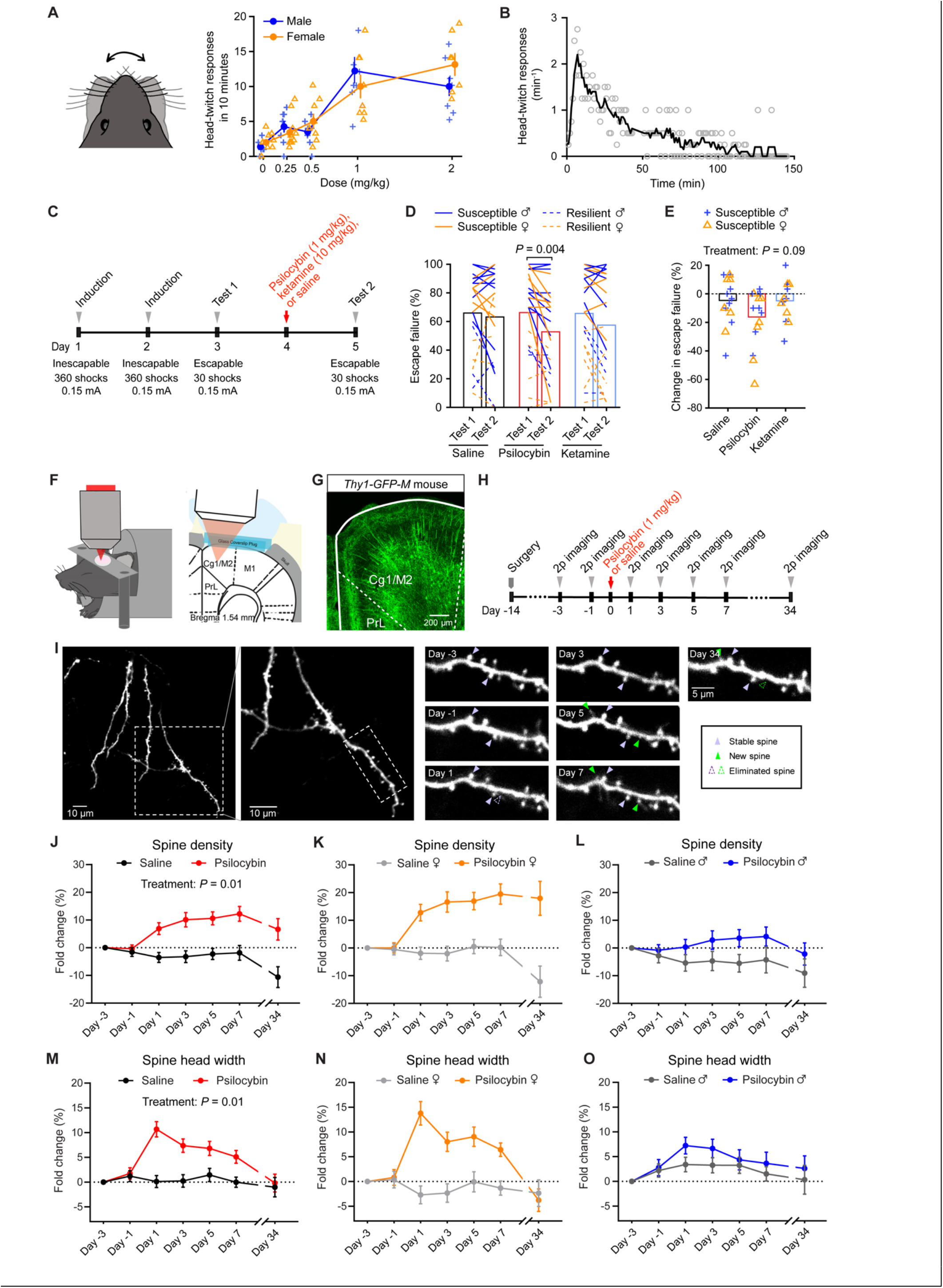
Psilocybin increases the density and size of dendritic spines in the mouse medial frontal cortex. **(A)** Head-twitch responses as a function of dose, tested on 82 C57BL/6J mice. **(B)** Time course of head-twitch responses after administrating psilocybin (1 mg/kg, i.p.), averaged from 2 males and 2 female C57BL/6J mice. Line, moving average. **(C)** Timeline for the learned helplessness assay. **(D)** The proportion of escape failure for all animals in Test 1 and Test 2, i.e., before and after psilocybin (1 mg/kg, i.p.), ketamine (10 mg/kg, i.p.), and saline administration. **(E)** The change in escape failure, from Test 1 to Test 2, for susceptible animals for psilocybin, ketamine, and saline treatments. **(F)** Imaging setup. **(G)** Fixed coronal section from *Thy1*^*GFP*^ mice. **(H)** Timeline for the longitudinal imaging study. **(I)** Example field of view. **(J)** Effects of psilocybin or saline treatment on spine density, plotted as fold-change from baseline value on Day -3. Mean ± SEM. **(K, L)** Similar to (J), plotted separately for females and males. **(M–O)** Similar to (J–L) for spine head width. Sample sizes and details of the statistical analyses are provided in Supplementary Table 1.

Next, to determine if this dose is associated with mitigation of stress-related phenotypes in mice, we tested the effect of 1 mg/kg psilocybin in a learned helplessness paradigm (Chourbaji et al., 2005). Mice received prolonged stress in the form of repeated, inescapable footshocks over 2 induction sessions, then were tested for active avoidance behavior with escapable footshocks 1 day before and 1 day after treatment (**Figure 1C**). Susceptible animals are mice in a learned helpless state characterized by reduced attempts to escape from the footshocks. We used 68 C57BL/6J mice to compare psilocybin (1 mg/kg, i.p.) against saline and ketamine (10 mg/kg, i.p.), which served as negative and positive controls (Wu et al., 2021). Within individuals, psilocybin reduced the proportion of escape failures (*P*=0.004, *post hoc* Bonferroni-corrected *t*-test; **Figure 1D**). For susceptible animals, the psilocybin group had a decrease or no change in escape failures in all but 1 of the 16 mice tested (94%; **Figure 1E; Supplementary Figure 1A**). Across individuals, comparison between saline, ketamine, and psilocybin did not reveal a main effect of treatment (*P*=0.09, two-way ANOVA; **Figure 1E**), presumably because of the across-subject variability in the behavioral responses. Nevertheless, overall, these results indicate that psilocybin can ameliorate maladaptive behavior induced by uncontrollable stress in mice.

In the body, psilocybin is dephosphorylated to psilocin, an agonist of 5-HT_2A_ receptors that are densely expressed in apical dendrites of layer 5 pyramidal neurons in the medial frontal cortex of primates and rodents (Aghajanian and Marek, 1997; Jakab and Goldman-Rakic, 1998; Willins et al., 1997). We therefore hypothesize that psilocybin may modify the dendritic architecture in the medial frontal cortex. We used chronic two-photon microscopy to track apical dendritic spines in the cingulate/premotor (Cg1/M2) region of the medial frontal cortex of *Thy1*^*GFP*^ mice (line M), in which a sparse subset of infragranular (layer 5 and 6) pyramidal neurons express GFP (Feng et al., 2000) (**Figures 1F and 1G**). We imaged before and after administering psilocybin (1 mg/kg, i.p.) or saline at 2-day intervals and then again ∼1 month later for a total of 7 imaging sessions (**Figures 1H and 1I**). In total, we tracked 1,820 dendritic spines on 161 branches from 12 animals including 6 males and 6 females. Spine morphology was analyzed blind to experimental conditions using standardized procedures (Holtmaat et al., 2009). We took advantage of the longitudinal data to normalize the change in spine density as fold-change in individual dendritic segments. For statistical analyses, we used a mixed-effects model, including treatment, sex, and days as factors, as well as all interaction terms. Variation within mouse and dendrite across days was accounted by including random effects terms for dendrites nested by mice. Our results indicate that a single dose of psilocybin induces a significant elevation in spine density (+7±2% on Day 1, +12±3% on Day 7; main effect of treatment, *P*=0.011, mixed-effects model; **Figure 1J–L**), increase in the width of spine heads (+11±2% on Day 1, and +5±1% on Day 7; main effect of treatment, *P*=0.013; **Figure 1M–O**), and higher spine protrusion length (**Supplementary Figure 1B–1D**). Details for all statistical tests including sample sizes are provided in **Supplementary Table 1**.

### Psilocybin elevates the formation rate of dendritic spines *in vivo*

Increased spine density could be due to higher formation rate, lower elimination rate, or both. To distinguish between the possibilities, we analyzed the same dendritic segments across adjacent imaging sessions to determine the turnover of dendritic spines. In females, the spine formation rate increased by 8±2% after psilocybin (absolute values for the formation rate: 7±1% on Day -1, 15±2% on Day 1; main effect of treatment, *P*=0.034, mixed-effects model; **Figures 2A and 2B**). Likewise, the spine formation rate was higher by 4±2% in males after psilocybin (absolute values for the formation rate: 6±1% on Day -1, 10±2% on Day 1). By contrast, there was no change in the elimination rate of spines (**Figure 2C**). The increase in spine formation rate was highest shortly after psilocybin administration and then diminished in subsequent days to return to baseline level and in equilibrium with the elimination rate. These data therefore support the view that the long-term increase in spine density is due to an initial boost of enhanced spine formation.

**Figure 2.**
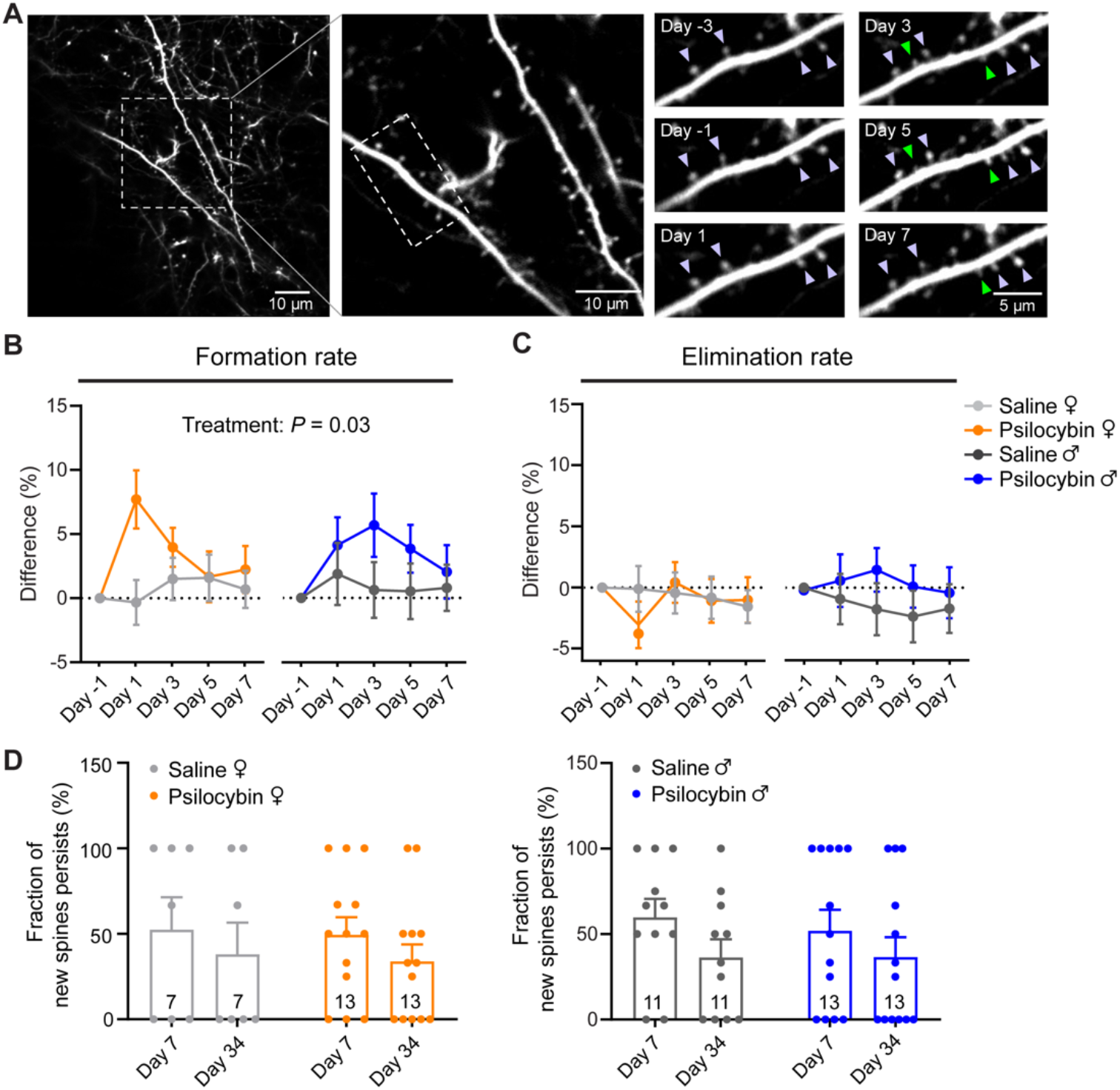
Psilocybin elevates the formation rate of dendritic spines. **(A)** Example field of view. Purple arrowhead, stable spine. Green arrowhead, new spine. **(B)** Effects of psilocybin or saline treatment on the formation rates of dendritic spines for female and male mice, plotted as difference from baseline value on Day -1. Mean ± SEM. **(C)** Similar to (B) for elimination rates. **(D)** Fraction of spines newly formed on Day 1 that remained stable on Day 7 and Day 34 for female and male mice. Filled circles, individual dendritic segments. Sample sizes and details of the statistical analyses are provided in Supplementary Table 1.

### A fraction of the psilocybin-induced spines is persistent for at least a month

A key question was whether the new spines formed after psilocybin administration would persist, because nascent dendritic spines can take 4 days to mature into functional synapses (Knott et al., 2006). For this reason, we tracked the new spines formed after psilocybin on Day 1 and found that about half of them remained stable on Day 7 (49±10% for females, 52±12% for males; **Figure 2D**). This suggests that a portion of the new dendritic spines induced by psilocybin would become functional synapses. Furthermore, because clinical trials indicated that psilocybin may provide long-term benefits for up to several months, for a subset of 4 mice, we imaged yet again at a further time point at 34 days after administration to find that a fraction of the psilocybin-evoked new spines remained persistent (34±10% for females, 37±12% for males; **Figure 2D and Supplementary Figure 2**). Psilocybin-induced spines were not significantly different, and therefore no less stable than spines formed in control conditions (main effect of treatment, *P*=0.9, two-way repeated-measures ANOVA). Intriguingly, select individual dendritic branches appeared to retain all the new spines, while other branches lost them almost completely, suggesting heterogeneity and potentially responsive and non-responsive subpopulations of pyramidal neurons. Altogether, these results demonstrate that a single dose of psilocybin induces rapid and long-lasting dendritic remodeling in layer 5 pyramidal neurons in the mouse medial frontal cortex.

### Ketanserin pretreatment, sufficient to abolish head-twitch responses, does not block psilocybin-induced structural plasticity

Multiple lines of evidence demonstrated that 5-HT_2A_ receptors are essential for serotonergic psychedelics’ psychotomimetic effects in humans (Vollenweider et al., 1998) and head-twitch responses in mice (Gonzalez-Maeso et al., 2007; Keiser et al., 2009). To study whether the effects of psilocybin on structural plasticity may involve 5-HT_2A_ receptors, we reduced the number of available 5-HT_2A_ receptors in the brain by pre-treating animals with the 5-HT_2A_ receptor antagonist ketanserin (1 mg/kg, i.p.), 10 minutes prior to the administration of psilocybin (1 mg/kg, i.p.) or saline. Behaviorally, the ketanserin pretreatment abolished completely the psilocybin-induced head-twitch responses (**Figure 3A**). Next, we repeated the two-photon imaging experiments in ketanserin-pretreated mice and tracked 1,443 dendritic spines on 120 branches from 8 animals including 4 males and 4 females (**Figure 3B**). We found that although the enhancing effect of psilocybin on spine density was no longer statistically significant (+5±2% on Day 1, +8±2% on Day 7; main effect of treatment, *P*=0.09, mixed-effects model; **Figure 3C; Supplementary Figure 3A and 3B**), there were still detectable increases in spine head width (+12±1% on Day 1, and +12±1% on Day 7; main effect of treatment, *P*=0.01; **Figure 3D; Supplementary Figure 3C and 3D**), spine protrusion length (**Supplementary Figure 3E– G**), and spine formation rate (absolute values for the formation rate: 5±1% on Day -1, 10±2% on Day 1 for female mice; 8±1% on Day -1, 14±2% on Day 1 for male mice; **Figure 3E; Supplementary Figure 3H**). It was previously determined that 1 mg/kg of ketanserin led to only a ∼30% blockade of 5-HT_2A_ receptors in the rat neocortex (Smith et al., 1995), likely due to limited transport into the brain for rodents (Syvanen et al., 2009). Therefore, in agreement with a recent study in the hippocampus (Hesselgrave et al., 2021), our results demonstrate that while a moderate knockdown of 5-HT_2A_ receptor function eliminates head-twitch responses, it is not sufficient to abolish the psilocybin-induced structural remodeling in mice.

**Figure 3.**
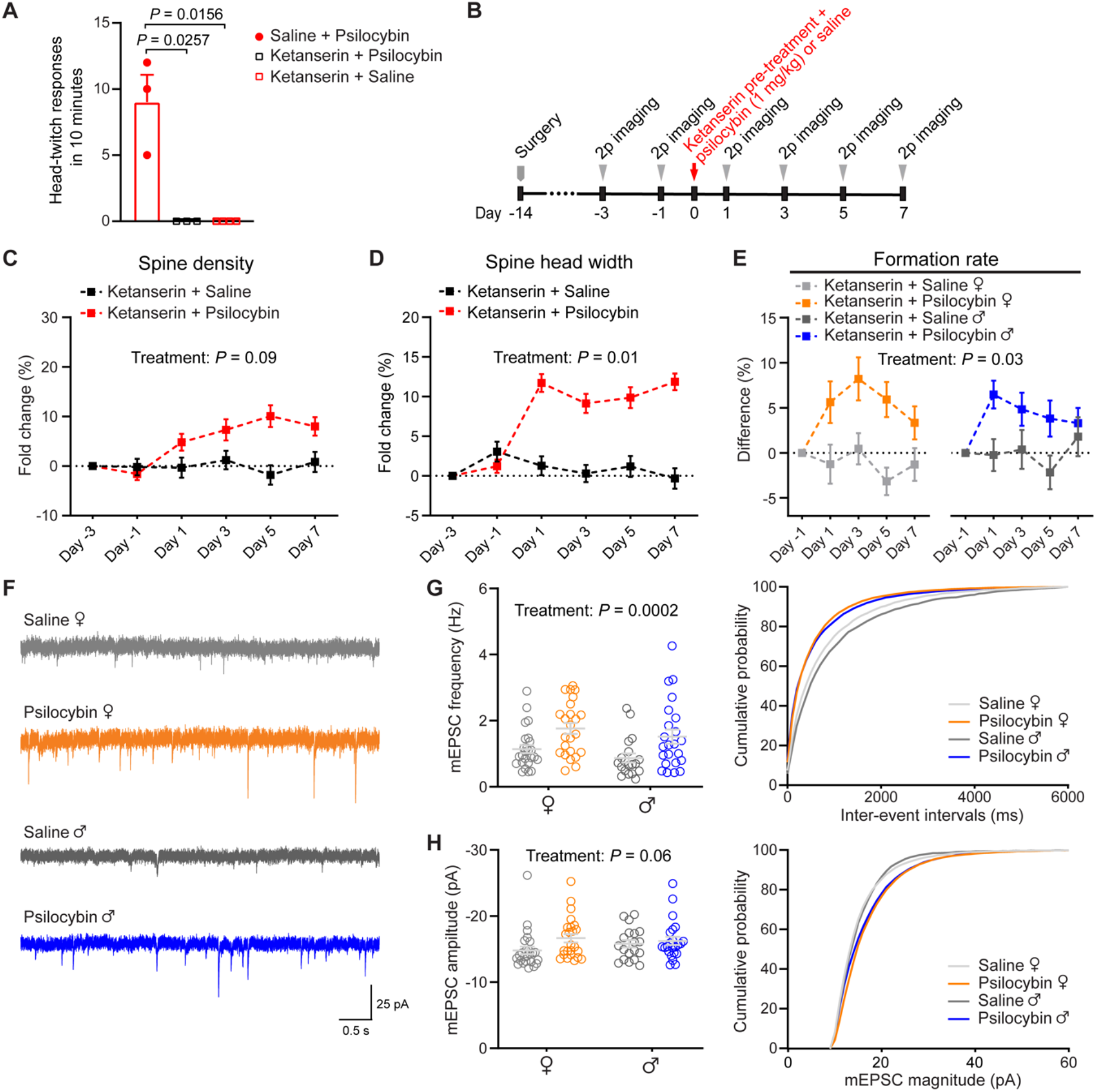
Mechanistic details revealed by ketanserin pretreatment and electrophysiological characterizations. **(A)** Head-twitch responses after administrating psilocybin without saline pre-treatment (10 min prior; n = 3 mice), psilocybin (1 mg/kg, i.p.) with ketanserin pre-treatment (1 mg/kg, 10 min prior; n = 3), and saline with ketanserin pre-treatment (n = 4). **(B)** Timeline for the experiment. **(C)** Effects of psilocybin or saline treatment on spine density in animals pretreated with ketanserin, plotted as fold-change from baseline value on Day -3. Mean ± SEM. **(D)** Similar to (C) for spine head width. **(E)** Effects of psilocybin or saline treatment on the formation rates of dendritic spines for female and male mice pretreated with ketanserin, plotted as difference from baseline value on Day -1. **(F)** Representative traces of mEPSCs recorded from putative layer 5 pyramidal neurons of Cg1/M2 in brain slices. **(G)** Grouped and cumulative distribution plots of mEPSC frequency for animals that received psilocybin or saline 24 h before recording. Each open circle denotes a cell (n = 25 cells from 4 females for saline; 24 cells from 4 females for psilocybin; 19 cells from 5 males for saline; 23 cells from 4 males for psilocybin). **(H)** Similar to (G) for mEPSC amplitude. Sample sizes and details of the statistical analyses are provided in Supplementary Table 1.

### Psilocybin elevates excitatory neurotransmission in medial frontal cortex

Most but not all dendritic spines are functional glutamatergic synapses. To elaborate on the effects of psilocybin on synaptic function, we performed whole-cell recordings in brain slices to measure miniature excitatory postsynaptic currents (mEPSCs) from putative layer 5 pyramidal neurons, identified based on morphology, in Cg1/M2 (**Figure 3F**). The results showed that, 24 h after treatment, we could detect an increase in mEPSC frequency in psilocybin-treated animals compared to saline controls (main effect of treatment, *P* = 0.0002, two-way ANOVA; **Figure 3G**). We also report a moderate effect of psilocybin on mEPSC amplitude (main effect of treatment, *P* = 0.06, two-way ANOVA; **Figure 3H**). Because mEPSC frequency and amplitude reflect the number and strength of synapses, these results demonstrate that the psilocybin-induced structural remodeling is accompanied by enhanced excitatory neurotransmission.

### Dependence of psilocybin-induced structural remodeling on brain region and dendrite type

To further support the conclusions, we tried to replicate the findings in a completely separate cohort of animals using a different approach. We administered *Thy1*^*GFP*^ mice with psilocybin (1 mg/kg, i.p.) or saline, sacrificed them 24 hours later, and imaged coronal brain sections using confocal microscopy. We expanded analyses to 6 areas of the brain, including 2 zones that encompass apical and basal dendrites and 3 regions of the frontal cortex: Cg1/M2, prelimbic/infralimbic (PrL/IL), and primary motor cortex (M1) (**Figures 4A–4C**). The results, consisting of 23,226 dendritic spines counted on 1,885 branches from 12 animals including 6 males and 6 females, reaffirmed the ability of psilocybin to promote the growth of new dendritic spines in Cg1/M2 in female mice (spine density: 0.46±0.02 versus 0.50 ±0.01 μm^-1^; **Figure 4D**). Effects of psilocybin on spine density were more pronounced in female animals than in male animals (treatment x sex, *P* = 0.013, two-way ANOVA; **Figure 4D**). We did not detect differences in spine protrusion length and spine head width (**Figures 4E and 4F**), which may be due to the across-subjects design, as we could not normalize the changes to the same dendritic branch and therefore this approach had less power than the within-subjects design of the chronic imaging experiment. We detected select morphological differences in PrL/IL and M1, including increases in spine protrusion length in PrL/IL (main effect of treatment, P = 0.026, two-way ANOVA), spine density in M1 in females (treatment x sex, *P* = 0.021, two-way ANOVA), and spine head width in M1 in females (treatment x sex, *P* = 0.008, two-way ANOVA), suggesting that the plasticity-promoting impact may not be unique to Cg1/M2 (**Figures 4G–4N; Supplementary Figure 4**). Furthermore, psilocybin had significant impact on basal dendrites in Cg1/M2, leading to higher spine density and spine protrusion length (spine density: main effect of treatment, *P* = 0.0004; spine protrusion length: main effect of treatment, *P* = 0.012, two-way ANOVA, **Figures 4O–4R**). Overall, the two sets of data converge to show that psilocybin promotes the growth of dendritic spines in layer 5 pyramidal neurons in the medial frontal cortex.

**Figure 4.**
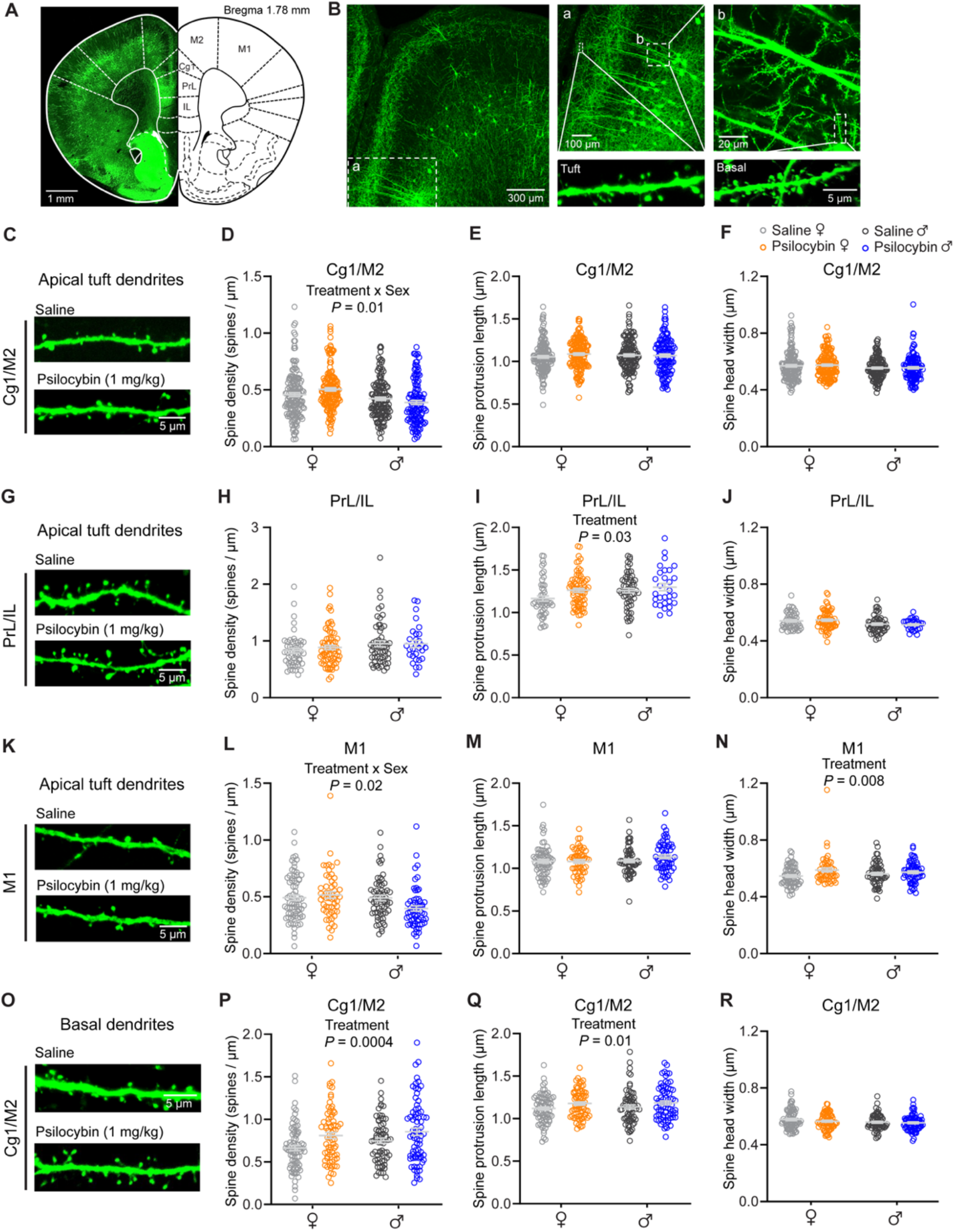
Region-specific effects of psilocybin. **(A)** Stitched confocal image of a coronal brain section from a *Thy1*^*GFP*^ mouse. **(B)** Magnified images showing apical and basal dendritic segments. **(C)** Images of apical dendrites in Cg1/M2. **(D)** Effects of psilocybin and saline on spine density for apical dendrites in Cg1/M2. Open circles, individual dendritic segments. Gray line, mean ± SEM. **(E)** Similar to (D) for spine protrusion length. **(F)** Similar to (D) for spine head width. **(G – J)** Similar to (C – F) for PrL/IL. **(K – N)** Similar to (C – F) for M1. **(O – R)** Similar to (C – F) for basal dendrites in Cg1/M2. Sample sizes and details of the ANOVA models are provided in Supplementary Table 1.

## Discussion

This study demonstrates that a single dose of psilocybin evokes growth of dendritic spines in the medial frontal cortex of the mouse. The persistence of the neural modifications is notable and may relate to the compound’s therapeutic effects for at least two reasons. First, depression is associated with a loss of synapses in the frontal cortex (Holmes et al., 2019). Restoring the number of neuronal connections may correct such deficit, providing a biological mechanism for alleviating symptoms of depression. Second, structural remodeling is integral to learning and facilitates the storage of lifelong memories (Xu et al., 2009; Yang et al., 2009). Psilocybin-induced neural plasticity could prime the brain for integrating new psychological experiences. Regardless of relative importance of these mechanisms, which are not mutually exclusive, our results indicate that the underlying structural trace in the brain is enduring and can be observed a long time after the initial drug exposure.

There is an ongoing debate over whether the hallucinogenic effects of serotonergic psychedelics are dissociable from the therapeutic effects (Olson, 2020; Yaden and Griffiths, 2020). Consistent with another new study (Hesselgrave et al., 2021), our results indicate that structural remodeling in the medial frontal cortex is undeterred by a moderate knockdown of 5-HT_2A_ receptor availability. The possibility to disrupt psilocybin’s acute behavioral effects without abolishing structural plasticity actions has clear implications for treatment in the clinic. However, it is not yet clear if the results will extrapolate to humans, because 5-HT_2A_ receptors have species-dependent differences in dissociation kinetics with serotonergic psychedelics (Kim et al., 2020). Moreover, our results do not rule out the involvement of 5-HT_2A_ receptors because this dose of ketanserin only blocks ∼30% of 5-HT_2A_ receptors in rodents (Smith et al., 1995), and the unaffected receptors might be enough to drive the dendritic remodeling. This number may be compared to the 50-70% 5-HT_2A_ receptor occupancy level required for the more intense psilocybin-induced psychological experience in humans (Madsen et al., 2019). Future studies with region- and cell-type-specific knockout of serotonin receptor subtypes are needed to produce more decisive evidence on the role of 5-HT_2A_ and other receptors in mediating the effects of psilocybin on dendritic plasticity.

By showing that the time course for psilocybin-induced structural remodeling is rapid and persistent *in vivo*, our study suggests that synaptic rewiring may be a mechanism shared by compounds with rapid antidepressant effects. Of note, the timing of psilocybin’s effect on the neural architecture is reminiscent of ketamine, which at subanesthetic dose causes similar rapid increase in spine density and elevation of spine formation rate in the medial frontal cortex (Moda-Sava et al., 2019; Phoumthipphavong et al., 2016). However, still unknown is how drugs with disparate molecular targets may yield comparable modifications on neural architecture and behavior (Kadriu et al., 2021; Savalia et al., 2021). Elucidating the mechanisms will be crucial towards unraveling the neurobiology of rapid-acting antidepressants.

## Acknowledgements

We thank Ben Kelmendi, Chris Pittenger, and Mikael Palner for discussions, Jane Taylor for use of open-field activity boxes, Huriye Atilgan and Heather Ortega for assistance on setting up video recording, and Adam Halberstadt for advice on scoring head-twitch response. Psilocybin was generously provided by Usona Institute’s Investigational Drug & Material Supply Program; the Usona Institute IDMSP is supported by Alexander Sherwood, Robert Kargbo, and Kristi Kaylo in Madison, WI. This work was supported by the Yale Center for Psychedelic Science, NIH/NINDS training grant T32NS041228 (C.L.), and NIH/NIGMS Medical Scientist Training grant T32GM007205 (N.K.S. and P.A.D.). We thank the Yale Center for Advanced Light Microscopy Facility for their assistance with confocal imaging, supported in part via NIH grant S10OD023598.

## Author Contributions

L.X.S. and A.C.K. designed the research. L.X.S. performed the two-photon and confocal imaging experiments and analyzed the two-photon imaging data. N.K.S. blinded and I.G. analyzed the confocal imaging data. C.L. performed and analyzed the behavioral experiments. I.G. and K.D. assisted with analyzing the behavioral data. P.A.D. performed the electrophysiological recordings. L.X.S. blinded and P.A.D. analyzed the electrophysiological data. N.K.S. assisted with the statistical analyses. L.X.S. and A.C.K. wrote the paper, with input from all other authors.

## Declaration of Interests

A.C.K. received psilocybin from the investigational drug supply program at Usona Institute, a non-profit organization. The authors declare no other competing interests.

## Methods

### Resource availability

#### Lead contact

Further information and requests for resources and reagents should be directed to and will be fulfilled by the Lead Contact Alex C. Kwan.

#### Materials availability

This study did not generate new unique reagents.

#### Data and Code Availability

The data that support the findings and the code used to analyze the data in this study will be made publicly available at https://github.com/Kwan-Lab.

### Experimental model and subject details

All experiments were performed on males and females. Animals were randomly assigned in the saline and psilocybin groups. No animals were excluded from data analysis. *Thy1*^*GFP*^ line M (Tg(Thy1-EGFP)MJrs/J, Stock No.007788) transgenic mice and C57BL/6J (Stock No. 000664) mice were obtained from Jackson Laboratory. For head-twitch response and learned helplessness, 6 to 10-week-old C57BL/6J mice were used. For electrophysiology, 6 to 8-week-old C57BL/6J mice were used. For two-photon imaging, *Thy1*^*GFP*^ mice underwent surgery when they were 6 to 8-week-old and then were used for imaging ∼2 weeks later. For confocal imaging, 8 to 12-week-old *Thy1*^*GFP*^ mice were used. Mice were group housed (2 – 5 mice per cage) under controlled temperature in a 12hr light–dark cycle (7:00 AM to 7:00 PM) with free access to food and water. Animal care and experimental procedures were approved by the Institutional Animal Care & Use Committee (IACUC) at Yale University.

### Method details

#### Psilocybin

Psilocybin was obtained from Usona Institute’s Investigational Drug & Material Supply Program. The chemical composition of psilocybin was confirmed by high performance liquid chromatography.

#### Head-twitch response

Head-twitch response was evaluated using 40 male and 42 female C57BL/6J mice. Upon arrival, animals habituated at the housing facility for >2 weeks before behavioral testing. Behavioral testing took place between 10:00 AM and 4:00 PM. Animals were weighed and injected intraperitoneally with saline or psilocybin (0.25, 0.5, 1, or 2 mg/kg). For ketanserin pretreated groups, animals received ketanserin (1 mg/kg, i.p.; S006, Sigma-Aldrich) 10 min before administration of saline or psilocybin (1 mg/kg, i.p.). Meanwhile, a group of animals received saline (10 mL/kg, i.p.) 10 min before administration of psilocybin (1 mg/kg, i.p.) as positive controls. We tried a higher dose of ketanserin (4 mg/kg, i.p.; n = 8 mice), however animals became visibly ill. We measured head-twitch response in groups of two animals: after injections, the two animals were immediately placed into separate chambers, made by inserting a plastic divider to halve an open-field-activity box (12” W x 6” H x 10” D). The box was within a sound attenuating cubicle with a built-in near-infrared light source and a white light source (interior: 28” W x 34” H x 22” D, Med Associates Inc.). Videos were recorded by a high-speed (213 fps), near-infrared camera (Genie Nano M1280, Teledyne Dalsa) mounted overhead above the open-field-activity box. Typical recordings were 30 minutes long and, for a subset of mice (2 males and 2 females), extended to >150 minutes. Between each measurement, the open-field activity box was thoroughly cleaned with 70% ethanol. The videos were scored for head twitches by an experienced observer blind to the experimental conditions.

#### Learned helplessness

Learned helplessness was evaluated using 34 male and 34 female C57BL/6J mice. Upon arrival, animals habituated at the housing facility for >2 weeks before behavioral testing. Behavioral testing took place between 7:00 AM and 4:00 PM. All animals underwent a 5-day protocol, adapted from previously described procedures for mice (Chourbaji et al., 2005). An animal was placed in a shuttle box separated by a guillotine door which, when open, allowed the animal to shuttle between two compartments (16” x 6.5” x 8.5”, Med Associates Inc.). On Day 1, the mouse received an induction session which involved 360 inescapable footshocks (0.15 mA) with variable duration (1–3 s) and variable inter-shock interval (1–15 s). The guillotine door was open throughout the induction session. On Day 2, the animal received another induction session with the same parameters. On Day 3, the animal underwent Test 1. The test session began with the guillotine door opening. Each test session involved a series of 30 footshocks (0.15 mA). The animal would receive a footshock from the grid floor of the compartment it was presently in. Footshock was terminated if the animal shuttled to the other compartment (“escape”) or at the end of the 10 s if it did not shuttle (“escape failure”), whichever occurred earlier. Each footshock was followed by an inter-shock interval (30 s), during which the guillotine door was closed. Escape latency was defined as the time elapsed from onset of footshock to time crossing the guillotine door, measured by infrared photobeams, for escape trials. Escape latency was set to 10 s for escape failure trials. On Day 4, 24 hr after Test 1, the animal was weighed and injected with saline (10 mL/kg, i.p.), ketamine (10 mg/kg, i.p.), or psilocybin (1 mg/kg, i.p.) and then immediately returned to their home cages. On Day 5, 24 hr after treatment, the animal underwent Test 2 which followed the same procedures as Test 1. At the end, data from all animals were collated, and mice were classified as “resilient / non-learned helpless” or “susceptible / learned helpless” based on their performance in Test 1. Escape failures and escape latencies were used as indicators of learned helplessness, and a k-means (k = 2) clustering algorithm was applied for classification.

#### Surgery

Prior to surgery, the mouse was injected with carprofen (5 mg/kg, s.c.; 024751, Henry Schein Animal Health,) and dexamethasone (3 mg/kg, i.m.; 002459, Henry Schein Animal Health). During surgery, the mouse was anesthetized with isoflurane (3 – 4% for induction and 1 – 1.5% for the remainder of surgery) and fixed in a stereotaxic apparatus (David Kopf Instruments). The body of the mouse rested on a water-circulating heating pad (Stryker Corp) set to 38 °C. Petrolatum ophthalmic ointment (Dechra) was used to cover the animal’s eyes. The hair on the head was shaved, and the scalp was wiped and disinfected with ethanol pad and betadine. An incision was made to remove the skin and the connective tissue above the skull was removed. Subsequently, a dental drill was used to make a ∼3-mm-diameter circular craniotomy above the right medial frontal cortex (center position: +1.5 mm anterior-posterior, AP; +0.4 mm medial-lateral, ML; relative to bregma). Artificial cerebrospinal fluid (ACSF, containing (in mM): 135 NaCl, 5 HEPES, 5 KCl, 1.8 CaCl2, 1 MgCl2; pH 7.3) was used to irrigate the exposed dura above brain. A two-layer glass window was made from two round 3-mm-diameter, #1 thickness glass coverslip (64-0720 (CS-3R), Warner Instruments), bonded by UV-curing optical adhesive (NOA 61, Norland Products). The glass window was carefully placed over the craniotomy and, while maintaining a slight pressure, adhesive (Henkel Loctite 454) was used to secure the glass window to the surrounding skull. A stainless steel headplate was affixed on the skull with C&B Metabond (Parkell) centered on the glass window. Carprofen (5 mg/kg, s.c.) was given to the mouse immediately after surgery and on each of the following 3 days. The mouse would recover for at least 10 days after the surgery before the start of imaging experiments.

#### Two-photon imaging

The two-photon microscope (Movable Objective Microscope, Sutter Instrument) was controlled by ScanImage 2019 software (Pologruto et al., 2003). The laser excitation was provided by a tunable Ti:Sapphire femtosecond laser (Chameleon Ultra II, Coherent) and focused onto the mouse brain with a water-immersion 20X objective (XLUMPLFLN, 20x/0.95 N.A., Olympus). The laser power measured at the objective was ≤ 40 mW. To image GFP-expressing dendrites, the laser excitation wavelength was set at 920 nm, and a 475 – 550 nm bandpass filter was used to collect the fluorescence emission. During an imaging session, the mouse was head fixed and anesthetized with 1 – 1.5% isoflurane. Body temperature was controlled using a heating pad and DC Temperature Controller (40-90-8D, FHC) with rectal thermistor probe feedback. Each imaging session did not exceed 2 hours. We imaged apical tuft dendrites at 0 – 200 µm below the dura. To target Cg1/M2 region, we imaged within 0 – 400 µm of the midline as demarcated by the sagittal sinus. Multiple fields of view were imaged in the same mouse. For each field of view, 10 – 40-µm-thick image stacks were collected at 1 µm steps and at 1024 × 1024 pixels at 0.11 µm per pixel resolution. We kept the same set of imaging parameters for the different imaging sessions.

For longitudinal imaging, we would return to the same fields of view across imaging sessions by locating and triangulating from a landmark on the left edge of the glass window. Each mouse was imaged on days -3, -1, 1, 3, 5 and 7 relative to the day of treatment. A subset of mice (2 males and 2 females) was imaged additionally on day 34. On the day of treatment (day 0), there was no imaging, and the mouse was injected with either psilocybin (1 mg/kg, i.p.) or saline (10 mL/kg, i.p.). For ketanserin pretreated groups, animals received ketanserin (1 mg/kg, i.p.) 10 min before administration of psilocybin (1 mg/kg, i.p.) or saline (10 mL/kg, i.p.). After injection, the mouse was placed in a clean cage under normal room lighting to visually inspect for head-twitch responses for 10 minutes, before returning the mouse to its home cage.

#### Confocal imaging

Each mouse was injected with either psilocybin (1 mg/kg, i.p.) or saline (10 mL/kg, i.p.). At 24 hr after injection, the mouse was deeply anesthetized with isoflurane and transcardially perfused with phosphate buffered saline (PBS, P4417, Sigma-Aldrich) followed by paraformaldehyde (PFA, 4% in PBS). The brains were fixed in 4% PFA for 24 hours at 4 °C, and then 50-µm-thick coronal brain slices were sectioned using a vibratome (VT1000S, Leica) and placed on slides with coverslip with mounting medium. The brain slices were imaged with a confocal microscope (LSM 880, Zeiss) equipped with a Plan-Apochromat 63x/1.40 N.A. oil objective for dendritic spine imaging and a Plan-Apochromat 20x/0.8 N.A. objective for stitching images of an entire brain slice.

#### Brain slice preparation

Female and male mice were randomly selected to receive either psilocybin (1 mg/kg, i.p.) or saline (10 mL/kg, i.p.) 24 hours before the experiment. The experimenter performing the electrophysiological recordings and analysis was blinded to the treatment condition. Coronal brain slices containing Cg1/M2 were prepared following procedures in a prior study (Ali et al., 2020b). Briefly, mice were deeply anesthetized with isoflurane and rapidly decapitated. The brain was quickly isolated into ice-cold slicing solution containing (in mM): 110 choline, 25 NaHCO_3_, 11.6 sodium ascorbate, 7 MgCl_2_, 3.1 sodium pyruvate, 2.5 KCl, 1.25 NaH_2_PO_4_, 0.5 CaCl_2_, and 20 glucose. Acute coronal slices (300 μm thick) were cut with a vibratome (VT1000 S, Leica Biosystems). The vibratome chamber was surrounded by ice and filled with oxygenated slicing solution. Slices were incubated in artificial cerebral spinal fluid (aCSF) containing (in mM): 127 NaCl, 25 NaHCO_3_, 2.5 KCl, 2 CaCl_2_, 1.25 NaH_2_PO_4_, 1 MgCl_2_, and 20 glucose for 30 min at 34°C. The slices were then maintained at room temperature for a minimum of 30 min before recording. The slicing solution and aCSF were prepared with deionized water (18.2 MΩ-cm), filtered (0.22 μm), and bubbled with 95% O_2_ and 5% CO_2_ for at least 15 min prior to use and throughout the slice preparation and recording.

#### Whole-cell recording

Slices were placed into an open bath chamber and perfused constantly with aCSF (2-3 mL/min) supplemented with tetrodotoxin (0.5 μM; Abcam) and picrotoxin (50 μM) to block Na^+^ currents and GABA_A_ receptors for isolating miniature excitatory post-synaptic currents (mEPSCs). aCSF was warmed and maintained at 34°C via an inline heater with closed-loop feedback control. Recording pipettes were pulled from borosilicate glass (BF-150-86-10, Sutter Instruments) to a resistance of 2-4 MΩ with a puller (P97, Sutter Instruments) and filled with double-filtered (0.22 μm) internal solution containing (in mM): 100 CsMeSO_4_, 25.5 CsCl, 10 Glucose, 10 HEPES, 8 NaCl, 4 Mg-ATP, 0.3 Na_3_-GTP, and 0.25 EGTA (pH 7.3, adjusted with 1M CsOH). Liquid junction potential was calculated to be 12.1mV and was not corrected for in recordings. Slices were visualized using differential interference contrast in a microscope (BX51W, Olympus) with a CCD camera (Retiga Electro, QImaging). Putative layer 5 pyramidal neurons were targeted for recording based on morphological features including large cell body, prominent apical dendrite, and distance from the pia. Cells were targeted with a depth of at least 30 μm below the surface of the slice. Electrophysiological recordings were performed on neurons that initially formed a stable seal and subsequently broke in successfully to the whole-cell configuration. Recordings were amplified (MultiClamp 700B, Molecular Devices) and digitized at 20 kHz (Digidata 1550, Molecular Devices). Neurons were held at −70 mV during recording. Recordings were excluded if the holding current >200 pA when held at -70 mV or if the access resistance increased by >10% from baseline during the recording o if the access resistance exceeds 25 MΩ at any point of the recording. Analysis of mEPSCs was conducted offline using the Easy Electrophysiology software (Easy Electrophysiology Ltd), with a template search algorithm. All drugs and regents were obtained from Sigma-Aldrich or Tocris unless otherwise noted.

### Quantification and statistical analysis

#### Analysis of the imaging data

Analyses of the two-photon and confocal imaging data were mostly similar, with an additional pre-processing step for motion correction of the two-photon imaging data using the StackReg plug-in (Thevenaz et al., 1998) in ImageJ (Schneider et al., 2012). Structural parameters such as spine head width and spine protrusion length were quantified based on a standardized protocol (Holtmaat et al., 2009; Phoumthipphavong et al., 2016). Briefly, if a protrusion extended for >0.4 µm from the dendritic shaft, a dendritic spine was counted. The head width of a dendritic spine was measured as the width at the widest part of the head of the spine. The protrusion length of a dendritic spine referred to the distance from its root at the shaft to the tip of the head. The line segment tool in ImageJ was used to measure the distances. Change in spine density, spine head width and spine protrusion length across imaging sessions were shown as fold-change from the value measured on the first imaging session (day -3) for each dendritic segment. The spine formation rate was calculated as the number of dendritic spines newly formed between two consecutive imaging sessions divided by the total number of dendritic spines observed in the first imaging session. The spine elimination rate was calculated as the number of dendritic spines lost between two consecutive imaging sessions divided by the total number of dendritic spines observed in the first imaging session. To assess the long-term dynamics of the spine formation and elimination rates across imaging sessions, we calculated the difference from the baseline rate, which was the spine formation or elimination rate of the same dendritic segment before psilocybin and saline injection (i.e., from day -3 to day -1). To quantify the persistence of newly formed spines, we calculated the number of dendritic spines newly formed on day 1 that are still present on day 7 and day 34, and divided by the total number of newly formed dendritic spines on day 1.

#### Statistics

Sample sizes and statistical analyses for each experiment are listed in **Supplementary Table 1**. Sample sizes were selected based on previous experiments reported in related publications (Grutzendler et al., 2002; Phoumthipphavong et al., 2016). GraphPad Prism 8 and R were used for statistical analysis. In the figures, data are presented as the mean ± SEM per dendritic branch.

For learned helplessness, we used a mixed effects model to test how proportion of escape failures (dependent variable) was impacted by fixed effects of treatment (saline vs. ketamine vs. psilocybin), test number (Test 1 vs. Test 2), and sex (female vs. male), including all second and higher-order interaction terms. Within-mouse variation was included as a random effects term. *Post hoc* paired-samples *t*-tests were used to analyze the change in Day 1 and Day 2 proportion of escape failures for the three treatment conditions, using Bonferroni correction for multiple comparisons. For ketanserin pretreatment experiments, a Kruskal-Wallis test (non-parametric one-way ANOVA) was used to test the difference in 10-minute head twitch responses across treatment groups (Saline + Psilocybin vs. Ketanserin + Psilocybin vs. Ketanserin + Saline), followed by Dunn’s multiple comparisons test for *post hoc* pairwise comparisons.

For *in vivo* two-photon imaging, dendritic spine scoring was performed while blind to treatment and time. Longitudinal measurements of dendrite structure were analyzed with mixed effects models for repeated measures using the *lme4* package in R. Linear mixed effects models were preferred to the commonly used repeated measures analysis of variance (ANOVA) due to fewer assumptions being made about the underlying data (e.g., balanced sampling, compound symmetry). Separate mixed effects models were created for each of five dependent variables: fold-change in spine density, fold-change in average spine head width, fold-change in average spine protrusion length, spine formation rate, and spine elimination rate. Each model included fixed effects for treatment (psilocybin vs. saline), sex (female vs. male), and time (Day 1, 3, 5, and 7) as factors, including all second and higher-order interactions between terms. Importantly, variation within mouse and dendrite across days was accounted by including random effects terms for dendrites nested by mice. Visual inspection of residual plots revealed no deviations from homoscedasticity or normality. *P*-values were calculated by likelihood ratio tests of the full model with the effect in question against the model without the effect in question. *Post hoc t*-tests were used to contrast psilocybin and saline groups per day, with and without splitting the sample by sex, applying Bonferroni correction for multiple comparisons. Spine persistence from two-photon imaging was analyzed with separate repeated measures ANOVAs for male and female mice, using fixed effects of treatment (psilocybin vs. saline), time (day 7 vs. day 34), and their interaction as independent predictors within dendrite. The same statistical analysis was applied to two-photon imaging data following ketanserin pretreatment, where the treatment groups were ketanserin + psilocybin vs. ketanserin + saline.

For electrophysiology data, blinding procedures involved one person inject psilocybin or saline, another person performing recording and measurements blind to treatment. Data were unblinded after all the measurements were completed. Two-way ANOVAs were used for mEPSC frequency and amplitude statistics. Treatment (psilocybin vs. saline), sex (female vs. male), and their interaction were included as independent predictors. *Post hoc t*-tests were used to contrast psilocybin and saline groups within sex, applying Bonferroni correction for multiple comparisons.

For confocal imaging data, blinding procedures involved one person performing imaging, another person scrambling the image file names, and a third person performing dendritic structural measurements blind to sex, treatment, and brain region. Data were unblinded after all of the measurements were completed. For each brain region in the confocal dataset (Cg1/M2, PrL/IL, and M1), separate two-way ANOVAs were constructed for apical and basal dendrites using spine density, spine head width, or spine protrusion length as the dependent variable. Treatment (psilocybin vs. saline), sex (female vs. male), and their interaction were included as independent predictors. *Post hoc t*-tests were used to contrast psilocybin and saline groups within sex, applying Bonferroni correction for multiple comparisons.

**Supplementary Video 1: Head-twitch response recorded with a high-speed camera; Related to Figure**

**1**. The mouse on the left received psilocybin (1 mg/kg, i.p.). The mouse on the right received saline. Video was recorded at 180 frames per second and played back at 30 frames per second, i.e., slowed down by 6 times. The left mouse had one head-twitch response during the video.

**Supplementary Table1. Detailed statistical analysis of all datasets, related to Figures 1, 2, 3, 4, and Supplementary Figures S1 to S4**.

---

**Supplementary Figure 1.**
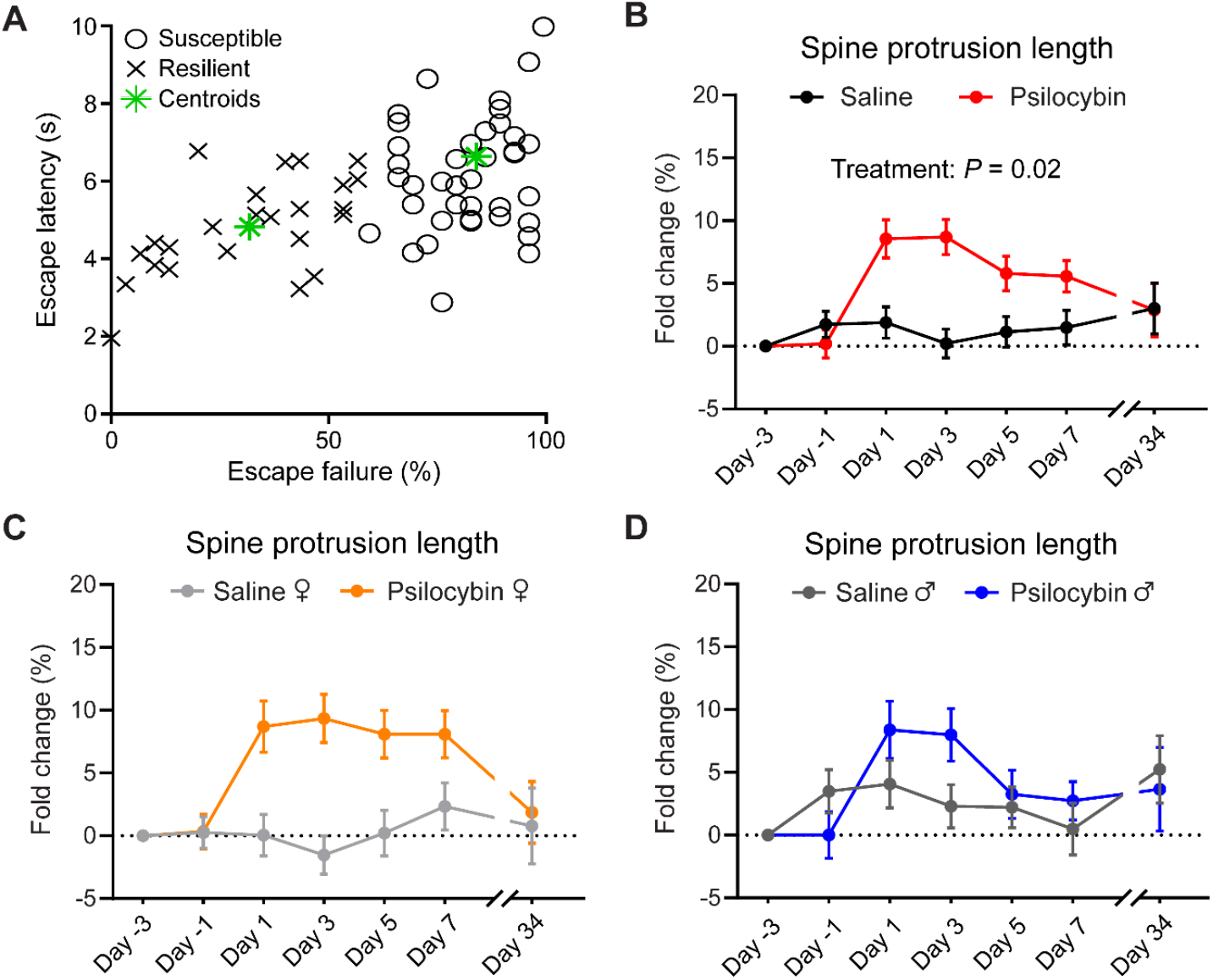
Details for learned helplessness and effects of psilocybin on spine protrusion length in Cg1/M2; Related to Figure 1. **(A)** Escape failure and escape latency in Test 1 for all the mice used in the learned helplessness assay (n = 68). A k-means clustering procedure was used to classify animals into resilient and susceptible groups. Green asterisk, the centroid location of each group. **(B)** Effects of psilocybin or saline treatment on spine protrusion length in *Thy1*^*GFP*^ mice, plotted as fold-change from baseline value on Day -3. **(C, D)** Similar to (B), plotted separately for females and males. Mean ± SEM. Sample sizes and statistical analyses are provided in Supplementary Table 1.

**Supplementary Figure 2.**
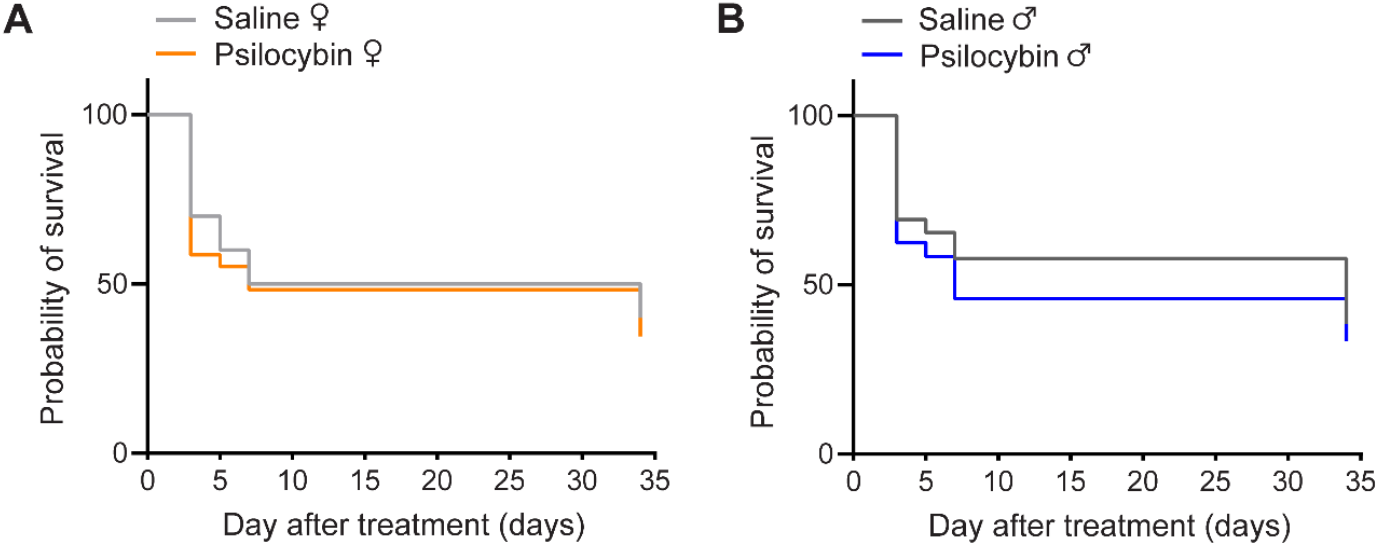
Survival curves for spines newly formed on Day 1; Related to Figure 2. **(A)** Survival curves of spines newly formed on Day 1 that remained stable in the following imaging sessions for female mice. **(B)** Similar to (A) for male mice. Mean ± SEM. Sample sizes and details of the ANOVA models are provided in Supplementary Table 1.

**Supplementary Figure 3:**
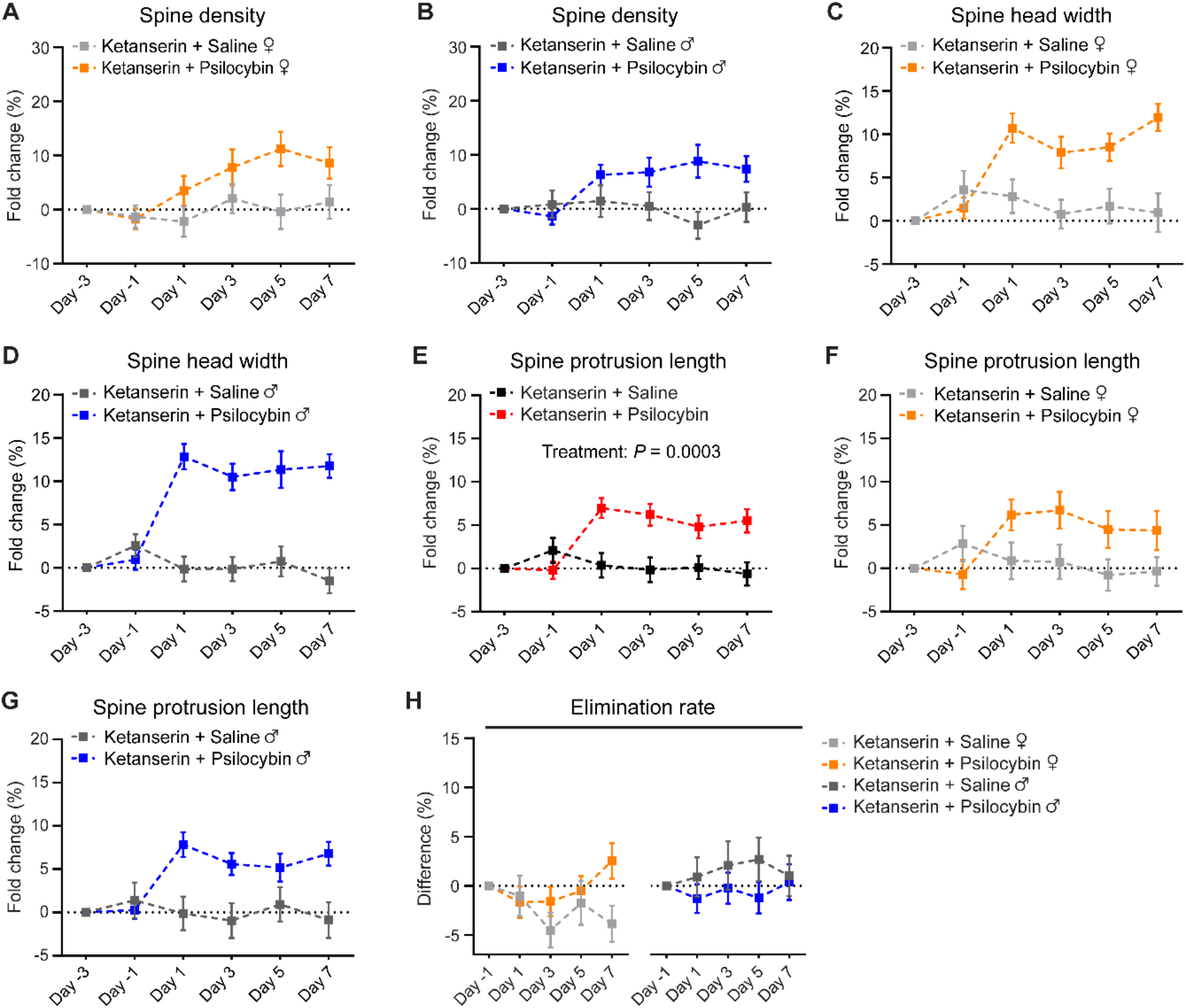
Psilocybin increases spine protrusion length in Cg1/M2 in mice with ketanserin pretreatment; Related to Figure 3. **(A, B)** Effects of psilocybin or saline treatment on spine density in animals pretreated with ketanserin, plotted as fold-change from baseline value on Day -3, plotted separately for females and males. Mean ± SEM. **(C, D)** Similar to (A, B) for spine head width. **(E)** Effects of psilocybin or saline treatment on spine protrusion length in *Thy1*^*GFP*^ mice with ketanserin pretreatment, plotted as fold-change from baseline value on Day -3. **(F, G)** Similar to (E), plotted separately for females and males. **(H)** Effects of psilocybin or saline treatment on the elimination rates of dendritic spines for female and male mice pretreated with ketanserin, plotted as difference from baseline value on Day -1. Sample sizes and statistical analyses are provided in Supplementary Table 1.

**Supplementary Figure 4.**
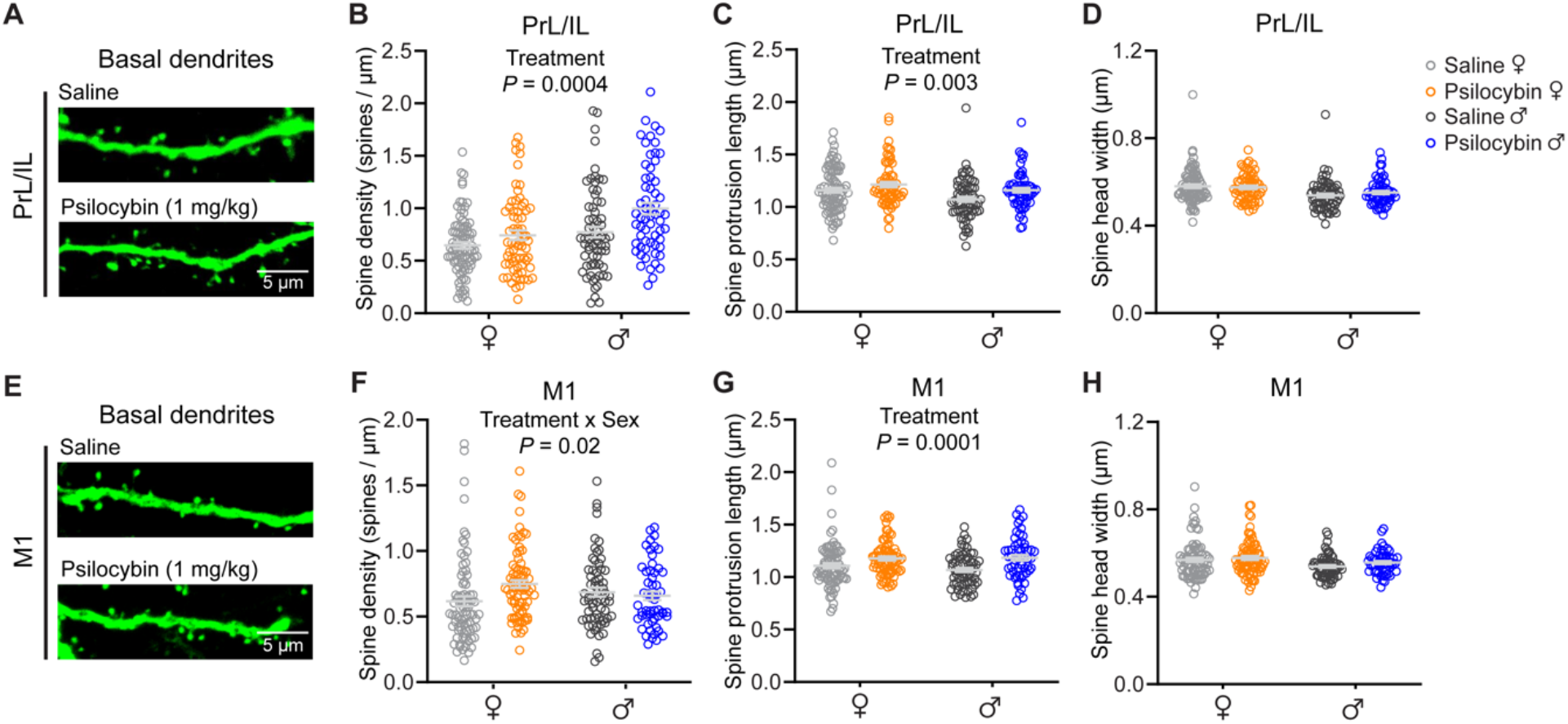
Effects of psilocybin on basal dendrites in PrL/IL and M1; Related to Figure 4. **(A)** Images of basal dendritic segments in PrL/IL from coronal brain sections from a *Thy1*^*GFP*^ mouse. **(B)** Effects of psilocybin and saline on spine density for basal dendrites in PrL/IL. Open circles, individual dendritic segments. **(C)** Similar to (B) for spine protrusion length. **(D)** Similar to (B) for spine head width. **(E – H)** Similar to (A – D) for M1. Mean ± SEM. Sample sizes and details of the ANOVA models are provided in Supplementary Table 1.

## Notes

### Summary of Updates

Added new data and revisions to the text.

